# Mapping In Vivo O-Glycoproteome Using Site-specific Extraction of O-linked glycopeptides (EXoO)

**DOI:** 10.1101/368282

**Authors:** Weiming Yang, Minghui Ao, Yingwei Hu, Qing Kay Li, Hui Zhang

## Abstract

Protein glycosylation is one of the most abundant post-translational modifications. However, detailed analysis of *in vivo* O-linked glycosylation, a major type of protein glycosylation, has been severely impeded by the scarcity of suitable methodologies. Here, we present a chemoenzymatic method for the site-specific <u>ex</u>traction <u>o</u>f <u>O</u>-linked glycopeptides (EXoO), which enabled the unambiguous mapping of over 3,000 O-linked glycosylation sites and definition of their glycans on over 1,000 proteins in human kidney tissues, T cells and serum. This large-scale localization of O-linked glycosylation sites nearly doubles the number of previously identified sites, demonstrating that EXoO is the most effective method to-date for defining the site-specific O-linked glycoproteome in different types of sample. Detailed structural analysis of the sites identified revealed conserved motifs and topological orientations facing extracellular space, the cell surface, the lumen of the ER and the Golgi. EXoO was also able to reveal significant differences in the *in vivo* O-linked glycoproteome of tumor and normal kidney tissues pointing to its broader use in clinical diagnostics and therapeutics.

## Introduction

Protein glycosylation is arguably the most diverse and sophisticated form of protein modification which drastically escalates protein heterogeneity to facilitate functional plasticity^1^. Compared to N-linked glycosylation, the study of O-linked glycosylation has proved difficult due to the structural complexity of the glycans added and the technical challenges posed to definitively characterizing them^2^. The process of O-linked glycosylation attaches different O-linked glycans to Ser and Thr residues, or less commonly Tyr residues in the human proteome^3^. Among the different types of O-linked glycosylation seen, O-linked N-acetyl-galactosamine (O-GalNAc) addition is a major type^4^. Definitive characterization of O-linked glycoproteins requires quantitative analysis of O-linked glycosylation sites and their corresponding glycans. In contrast to N-linked glycosylation, where a consensus glycosylation motif has been identified, there is no consensus O-linked glycosylation motif for the amino acid residues surrounding the glycosylated Ser or Thr. The cellular machinery for O-linked glycosylation, located primarily in the Golgi apparatus, is believed to operate stochastically in response to changes in a wide range of both intrinsic and extrinsic factors^1^. As a consequence, O-linked glycosylation can exhibit high heterogeneity in different cells, tissues and diseases.

The enrichment of O-linked glycopeptides from complex biological samples is an essential prerequisite to definitive identification of the otherwise low abundance O-linked glycoproteome^5^. To this end, a range of different enrichment methodologies have been reported including the use of lectins^6^, ^7^, HILIC^8^, ^9^, hydrazide chemistry^10^, ^11^, metabolic labelling^5^, ^12^ and a gene-engineered cell system named ‘SimpleCell’^13^. However, all of these methodologies have deficiencies that mean that the limited accessibility to the *in vivo* O-linked glycoproteome they afford remains as a severe hindrance to the structural and functional study of *in vivo* O-linked glycoproteins. In addition to the current difficulties in enrichment of O-linked glycopeptides from complex mixtures, the precise pinpointing of O-linked glycosylation sites and their corresponding site-specific glycans has proved to be even more challenging. Mass spectrometry (MS) using electron transfer dissociation (ETD) for fragmentation has to-date been the prevailing analytical tool to localize O-linked glycosylation sites^14^. Caveats associated with site localization using ETD render the method to be inefficient in mapping sites^15^, ^16^. Alternatively, chemical deglycosylation such as β-elimination Michael addition can be used to substitute O-linked glycans for glycosylation site identification^17^. However, a variety of complications and non-specific substitutions at other posttranslational modifications or at unmodified Ser and Thr limits the performance of this alternative method when employed on complex samples^17^. In sharp contrast, the identification of N-linked glycosylation sites and N-linked glycans is highly efficient making use of a specific enzyme (PNGase F) and the consensus NXS/T motif (X ≠ Pro)^18^. PNGase F simultaneously liberates N-linked glycans and converts the Asn in the glycosylation site to Asp (deamination) thereby identifying the site^19^, ^20^. Unfortunately, there is no PNGase F-equivalent for releasing O-linked glycans and localization of O-linked glycosylation sites. Consequently, any new methodology capable of large-scale analysis of site-specific O-linked glycoproteome *in vivo* would be highly advantageous.

Here, we introduce a new chemoenzymatic method for <u>ex</u>traction <u>o</u>f site-specific <u>O</u>-linked glycopeptides, named EXoO. It has been designed to simultaneously enrich and identify *in vivo* O-linked glycosylation sites and define their site-specific glycans. EXoO comprises four steps:

i. digestion of protein samples to generate peptides, (ii) conjugation of peptides to a solid-support, (iii) release of intact O-linked glycopeptides at O-linked glycosylation sites using an O-linked glycan-dependent endo-protease named OpeRATOR, (iv) analysis of the released intact O-linked glycopeptides by LC-MS/MS. OpeRATOR, identified from the mucin degrading human intestinal bacterium *Akkermansia muciniphila*, recognizes O-linked glycans and cleaves O-linked glycopeptides at the N-termini of O-linked glycan-occupied Ser or Thr to release site-specific O-linked glycopeptides with the glycosylation sites at the N-terminus of the peptide for unambiguous localization. EXoO was developed using bovine fetuin and then benchmarked to identify a plethora of O-linked glycosylation sites using human kidney tissues, serum and T cells before being applied to defining the aberrant O-linked glycoproteome in human kidney tumor tissue.

## Results

### Extraction of site-specific O-linked glycopeptides

In EXoO, proteins are first digested to generate peptides, which are then conjugated to a solid-support. After washing, the O-linked glycopeptides are enzymatically released from the support using an endo-protease OpeRATOR that requires the presence of O-linked glycans to specifically cleave on the N-terminal side of O-linked glycan-occupied Ser or Thr (Fig. 1A). To demonstrate proof of principle, bovine fetuin was analysed and the six known O-linked glycosylation sites documented in the Uniprot database were pinpointed at Ser-271, Thr-280, Ser-282, Ser-296, Thr-334 and Ser-341 (Supplementary Table 1). In addition, a new O-linked glycosylation site at Ser-290 was also identified (Supplementary Table 1 and Supplementary Fig. 1). Of note, O-linked glycans were still attached to the site-specific O-linked glycopeptides as confirmed by the detection of oxonium-, peptide (Y0)- and less commonly peptide + HexNAc (Y1)-ions in the MS/MS spectrum (Fig. 1B). The detection of oxonium ions in the MS/MS spectrum is particularly useful for obtaining the correct identification of O-linked glycopeptides. In addition, the chemical conjugation of peptides to a solid-support allows efficient washing and specific enzymatic release of intact O-linked glycopeptides. As a result, 290 peptide spectrum matches (PSMs) were assigned to fetuin site-specific O-linked glycopeptides with Ser or Thr at the N-termini of peptides, glycan modification and oxonium ions in the MS/MS spectra from a total of 415 assigned PSMs, indicating a specificity of approximately 70% for O-linked glycopeptide enrichment using EXoO (Supplementary Table 1). The analysis of fetuin demonstrated the ability of EXoO to enrich and identify O-linked glycopeptides at specific O-linked glycosylation sites and their corresponding O-linked glycans.

**Figure 1.**
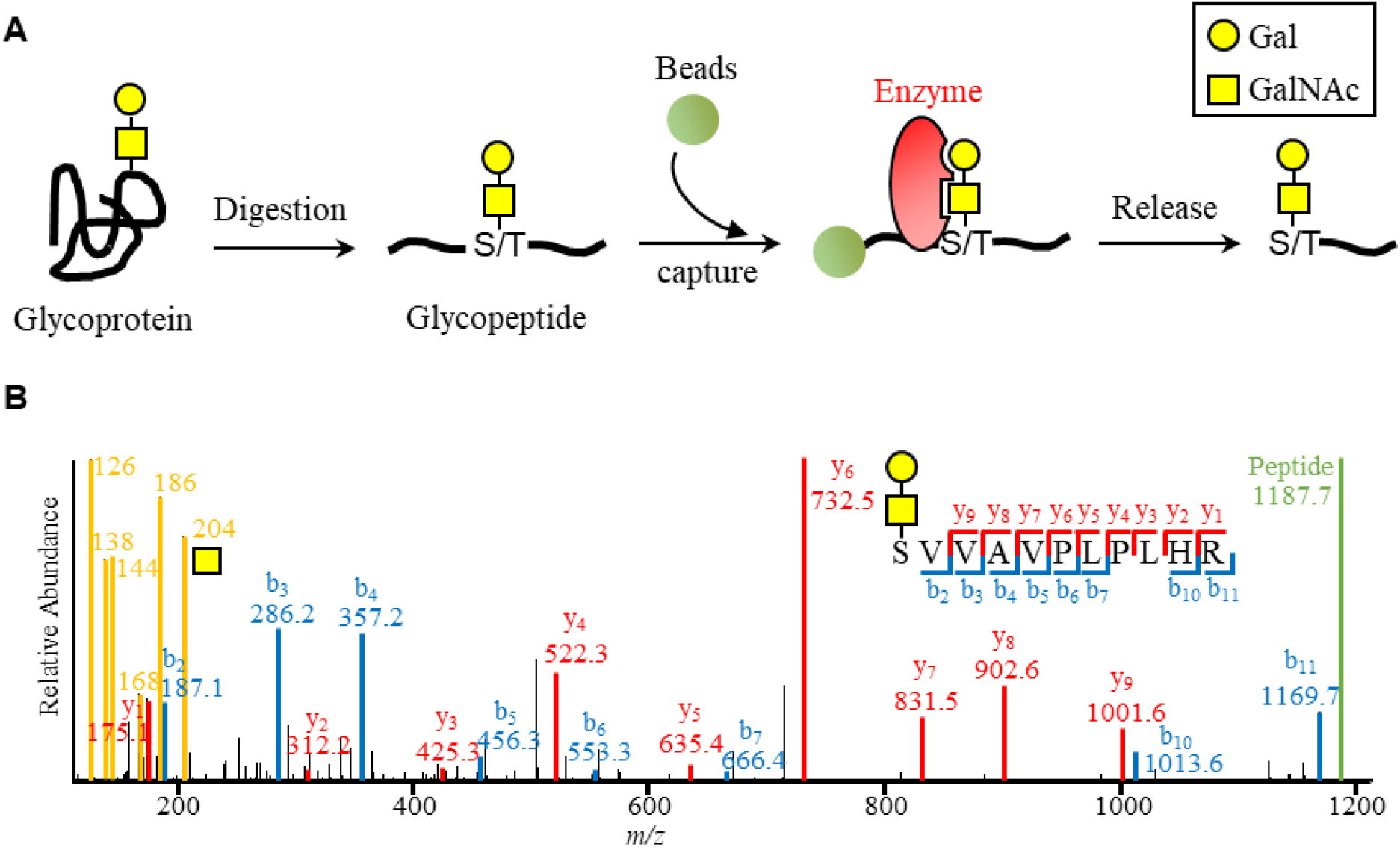
EXoO procedure for mapping the site-specific O-linked glycoproteome. (A) Schematic of EXoO process for precision mapping of O-linked glycosylation sites and site-specific glycans. (B) MS/MS spectrum of the site-specific O-linked glycopeptide at Ser-296 in bovine fetuin.

### Large-scale and precision mapping of the O-linked glycoproteome in human kidney tissue, T cells and serum

EXoO was benchmarked using human kidney tissue, T cells and serum to determine performance of the method in samples with differing levels of protein complexity. To do this, O-linked glycopeptides were extracted using EXoO and fractionated into 24 fractions and then subjected to LC-MS/MS analysis (Fig. 2A). To study kidney tissue, paired tumor and normal tissues were collected from three patients. The extracted proteins from these tissues were separately pooled to generate two samples i.e. tumor and normal. After analysis with 1% false discovery rate (FDR) at PSM level, 35,848 PSMs were assigned to 2,804 O-linked glycopeptides containing 1,781 O-linked glycosylation sites from 590 glycoproteins (Supplementary Table 2). When the EXoO approach was applied to the analysis of T cells, 4,623 PSMs were assigned to contain 1,295 O-linked glycosylation sites from 1,982 O-linked glycopeptides and 590 glycoproteins (Supplementary Table 3). Finally, human serum that contains a number of highly glycosylated proteins and has been previous subjected to detailed mapping of N-linked glycosylation sites and N-linked glycans but for which there has been little success in mapping of O-linked glycosylation sites and O-linked glycans^21^–^25^ was analyzed. With 1% FDR, 6,157 PSMs were assigned to 1,060 O-linked glycopeptides with 732 O-linked glycosylation sites from 304 glycoproteins being identified (Supplementary Table 4). This analysis of human tissue, T cells, and serum demonstrated that EXoO is highly effective tool for accessing the O-linked glycoproteome in different types of samples.

**Figure 2.**
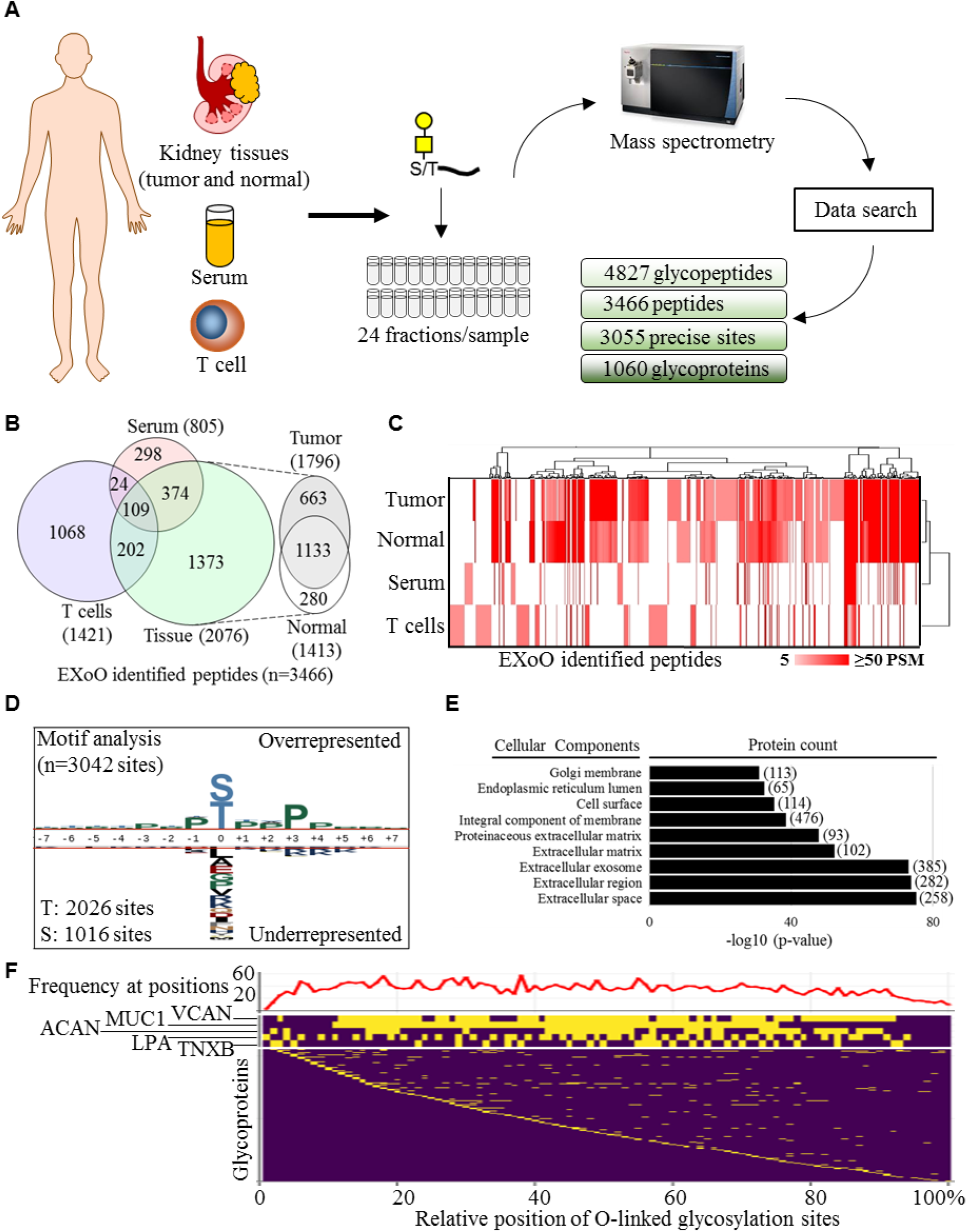
Mapping of *in vivo* O-linked glycoproteome. (A) Workflow to map site-specific O-linked glycoproteome in human samples (B) Distribution of EXoO identified peptides in different samples. (C) Unsupervised hierarchical clustering of samples and PSM number of peptides to show that EXoO identified peptides exhibited different distribution and relative abundance in samples. Euclidean distance and City-block distance were used for clustering of samples and peptides, respectively. (D) Analysis of amino acid sequence surrounding O-linked glycosylation sites. (E) Cellular component analysis of O-linked glycoproteins. (F) Landscape distribution of the O-linked glycosylation sites and its frequency in proteins. Relative position was calculated using amino acid position of the sites to divide total number of amino acids of the protein and time a hundred to show in percentage. Lower panel: relative position of the sites in proteins. Proteins with positions close to protein N-termini were ranked to the top. Middle panel: relative position of sites in five proteins with the highest number of sites. Protein with higher number of sites was ranked to the top. Upper panel: frequency of the sites in proteins. The y-axis of upper panel shows the number of sites at the relative position of proteins.

### Characterizing the *in vivo* O-linked glycoproteome

Our large-scale analysis unambiguously mapped 3,055 O-linked glycosylation sites from 1,060 glycoproteins in kidney tissues, T cells and serum (Supplementary Table 5). To compare the EXoO identified sites to that reported previously, 2,746 reported O-GalNAc sites were collected from O-GalNAc human SimpleCell glycoproteome DB^13^, ^26^, PhosphoSitePlus^27^ and Uniprot database^28^. Remarkably, EXoO identified 2,580 new O-linked glycosylation sites, an approximately 94% increase of the known sites, which however are mapped primarily using engineered cell lines.

To determine sample-specific O-linked glycoproteome, the distribution of EXoO identified peptides in different samples was determined. Kidney tissue and T cells had a large number of unique peptides compared to that seen for serum, with more than half of peptides detected in serum also being identified in the tissue sample, possibly due to the presence of serum in tissue samples (Fig. 2B). To visualize the relative abundance of peptides in different samples, the PSM numbers of peptides, which are suggestive of relative abundance, were clustered by unsupervised hierarchical clustering (Fig 2C). This showed that not only that the peptides differed between samples but also that their relative abundances were markedly divergent between samples (Fig. 2C). Interestingly, immunoglobulin heavy constant alpha 1 (IGHA1) has the highest PSM number in the normal tissue and serum but had the second highest PSM number in the tumor tissue where versican core protein (VCAN) scored the highest PSM number suggesting their relatively high abundance for detection and aberrant O-linked glycosylation of VCAN in tumor tissue. In the case of IGHA1, four of the five known sites on Ser residues and two new sites on Thr residues were mapped supportive of EXoO’s capacity to both localize known and discover new O-linked glycosylation sites. Overall, these data suggest that protein O-linked glycosylation is highly dynamic and may exhibit a disease-specific signature.

To identify possible O-linked glycosylation motifs, the amino acids (± 7 amino acids) at and surrounding 3,042 of the sites mapped in this study were analysed. O-linked glycan addition at Thr and Ser accounted for 67.6 and 22.4% of the sites, respectively (Fig. 2D). Analysis of the surrounding sequence motifs revealed that Pro was overrepresented at the +3 and –1 positions irrespective of which amino acid (Thr or Ser) was glycosylated or sample type (Fig. 2D and Supplementary Fig. 2). Overall enrichment of Pro was observed in the amino acids surrounding O-linked glycosylation sites (Supplementary Fig. 2). Thirteen O-linked glycosylation sites were not used in the motif analysis because they were located close to the termini of proteins concerned and consequently did not have enough surrounding amino acids to allow for full motif analysis.

Gene ontology (GO) analysis of EXoO identified glycoproteins was carried out and this showed that extracellular space, the cell surface, the ER lumen and the Golgi membrane were the major cellular components for O-linked glycoproteins (Fig. 2E). Analysis of biological process and molecular function suggested various activities and functionalities associated with O-linked glycoproteins, consistent with their important role in different aspects of biology (Supplementary Fig. 3). Specifically, extracellular matrix organization, cell adhesion and platelet degranulation were the biological processes most represented in the glycoproteins identified (Supplementary Fig. 3); whereas heparin binding, calcium ion binding and integrin binding were the top molecular functions identified (Supplementary Fig. 3).

To overview the positional distribution of the O-linked glycosylation sites identified, the relative position of the sites in the proteins was determined and arranged relative to the N-terminus of the glycoprotein in question (Fig. 2F lower panel). In addition, frequency of the sites at the relative position of proteins was calculated (Fig. 2F upper panel). It was found that the sites had relatively even distribution across the protein sequence but less frequent at protein termini (Fig. 2F upper panel). Strikingly, 20 proteins were seen to contain over 20 sites. Five proteins with the highest number of sites were zoomed for clear visualization in Figure 2F middle panel. These heavily glycosylated proteins appeared to show continuous clusters of many vincinal sites that nearly cover the whole proteins such as VCAN, mucin-1 (MUC1) and aggrecan core protein (ACAN). The cluster of sites could be relatively short while distributed evenly as seen in apolipoprotein (LPA) and Tenascin-X (TNXB). Among these heavily O-linked glycoproteins, VCAN contained the highest number of sites reaching 165 sites with distinct peptide sequences surrounding the sites whereas MUC1 contained 161 sites, the second highest, but composited from only six distinct sequence repeats. ACAN, LPA and TNXB were heavily O-linked glycosylated to have 82, 73 and 44 sites, respectively. Analysis of the site distribution on glycoproteins demonstrated advantage of EXoO to study heavily O-linked glycoproteins that is difficult to be analyzed by current analytical approach due to structural complexity and resistance to enzymatic digestion.

To determine localization of the sites to protein structures, protein topological and structural annotations were retrieved from Uniprot database and mapped to the EXoO identified sites. It was found that approximately 28.3 and 10.3% of the sites were predicted to localize in extracellular and luminal region, respectively (Supplementary Fig. 4). In contrast, only approximately 1.6% of the sites were predicted in cytoplasmic compartment (Supplementary Fig. 4). Approximately 5% of the sites were associated with Ser/Thr/Pro-rich region but weaker correlation of the sites to other protein structures including repeats, coiled coil, beta strand, helix, turn and signal peptides (Supplementary Fig. 4). Close to none correlation of the sites to intra-and transmembrane region of proteins. The structural correlation of the sites to extracellular, lumen and Ser/Thr/Pro-rich regions coincided with the location of O-linked glycoproteins to present on extracellular space, the cell surface, the ER and the Golgi lumen for various functionality.

### Mapping aberrant O-linked glycoproteome associated with human kidney tumor

To identify changes in the O-linked glycoproteome between normal and tumor kidney tissue, spectral counting label-free quantification of the EXoO identified peptides was used (Fig. 3A). This identified 56 O-linked glycoproteins as exhibiting significant change using scoring criteria of at least a two-fold change together with a difference of at least 10 PSMs between normal and tumor samples (Fig. 3A, Supplementary table 6). The most striking change observed was the dramatic increase of O-linked glycans, primarily in the core 1 structure Hex(1)HexNAc(1) across the 163 and 82 sites mapped in VCAN and ACAN respectively (Fig. 3C). For example, 35 PSMs were found at Thr-2983 of VCAN from tumor tissue but none in normal tissue samples (Fig. 3C red asterisk in upper panel). Similarly, 109 PSMs were detected at Thr-374 of ACAN from tumor tissue and only five in normal tissue (Fig. 3C red asterisk in lower panel). VCAN and ACAN are known proteoglycans that have long sugar chains in normal condition^29^. The extensive addition of short core 1 O-linked glycans to these two proteins would be expected to enhance their mucin-type properties such as resistance to enzymatic digestion and improved stiffness and this in turn may alter their biological and biomechanical properties producing some remodelling of the tumor microenvironment^30^. In addition to VCAN and ACAN, an average of 4.3-fold increase was detected in 14 sites across fibulin-2 (FBLN2), a glycoprotein known to be involved in stabilizing the VCAN and ACAN network for growth and metastasis of tumor^30^, ^31^ (Fig. 3D and Supplementary table 6). Remodelling of the extracellular matrix (ECM) in tumor tissue might be further underpinned by the significant changes of other ECM related proteins such as ELN, LTBP1, LTBP2, LTBP3, EMILIN2, FN1, CDH16, EMCN and collagens COL8A1, COL12A1, COL28A1, COL26A1 and enzymes including ITIH2, MMP14, ADAMTS7, SERPINE1, ANPEP, PAPLN, ADAMTSL5, GGT5 and CPXM1 detected in tumor tissue (Fig. 3D and Supplementary table 6). This type of fine-tuning of the ECM network is known to be critical to supporting tumorigenesis and tumor progression^31^. In addition, carbonic anhydrase 9 (CA9) and angiopoietin-related protein 4 (ANGPTL4), proteins known to respond to tumor hypoxia^32^, ^33^, showed 13 and 3.6-fold increases respectively in tumor tissue (Fig. 3D and Supplementary table 6). Finally, EGF-containing fibulin-like extracellular matrix protein 1 (EFEMP1) which binds to epidermal growth factor receptor (EGFR) to promote tumor growth, invasion and metastasis^34^, showed a 3-fold increase across seven O-linked glycosylation sites in tumor tissue (Fig. 3D and Supplementary table 6). By contrast, both IGHA1and MUC1, known O-linked glycoproteins, showed no detectable change between normal and tumor tissue indicating that the changes in O-linked glycoproteins observed in tumor tissue are highly selective.

**Figure 3.**
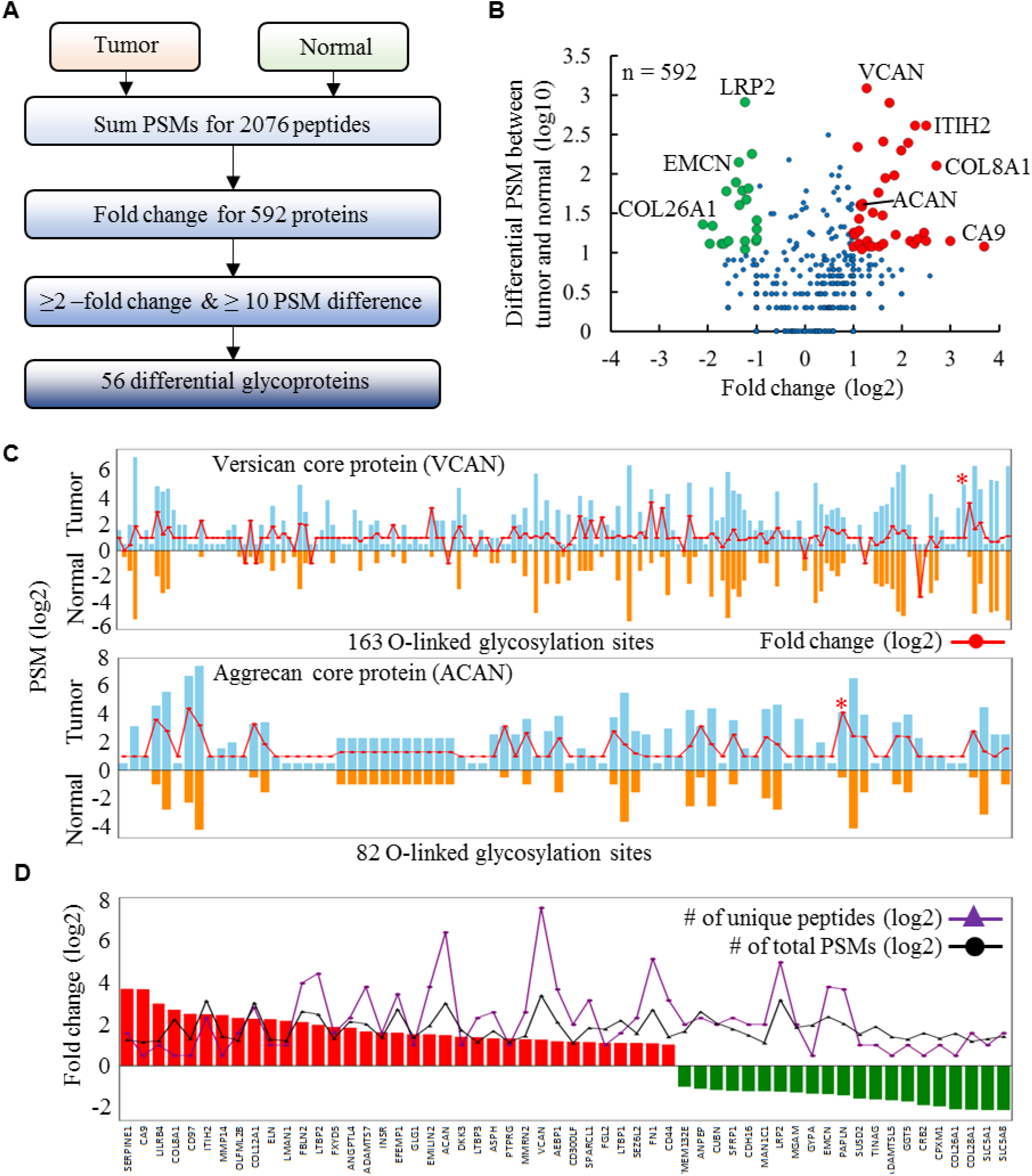
Comparative analysis of the O-linked glycoproteome of normal and tumor derived kidney tissue. (A) Data processing steps in label-free quantification to identify differential glycoproteins between tumor and normal kidney tissues. (B) Volcano plot of differentially expressed O-linked glycoproteins between tumor and normal tissues. (C) Extensive addition of O-linked glycans at sites covering versican core protein (VCAN) (upper panel) and aggrecan core protein (ACAN) (lower panel). Fold change for specific site between tumor and normal was showed as connected dots in red. Sites only detected in tumor or normal were assigned 2- or 0.5-fold change, respectively. Red asterisks indicate sites with the highest PSM number divergent between tumor and normal tissues. (D) The 56 O-linked glycoproteins with significant change between tumor and normal.

## Discussion

A novel tool (EXoO) has been developed for the combined mapping of O-linked glycosylation sites in proteins and the definition of the O-linked glycans at those sites. The main advantages of EXoO are (i) applicability for *in vivo* analysis; (ii) precise localization of O-linked glycosylation sites; (iii) simultaneous definition of O-linked glycans at the glycosylation sites; (iv) no requirement for ETD mass spectrometry to localize sites. The effectiveness of the method derives from the specific enrichment of O-linked glycopeptides at specific glycosylation sites using the tandem-action of a solid-support and the O-linked glycan specific OpeRATOR enzyme. The solid-support specifically binds peptides and maximizes the removal of non-bound molecules, while the OpeRATOR enzyme specifically cleaves on the N-terminal side of glycan-occupied Ser and Thr residues of the bound peptides to release O-linked glycosylation sites at the N-terminus of peptides enabling unambiguous localization of O-linked glycosylation sites. The O-linked glycan remains attached to the released O-linked glycopeptides and provides oxonium ions in the MS/MS spectrum to facilitate confident identification.

Analysis of the more than 3,000 O-linked glycosylation sites identified by EXoO revealed many glycoproteins that were previously not known to be modified by O-linked glycosylation. Many of those identified were mucin-type glycoproteins whose mucin domain contains clusters of dense O-linked glycans that protect the underlying peptide backbone from normal proteolytic digestion and consequently typical proteomic analysis would not contain detailed information on many of these domains. By contrast, OpeRATOR is naturally designed to dissect such mucin-type O-glycan-rich regions. Therefore, using EXoO allowed detailed mapping of over one hundred sites on VCAN and MUC1. Motif analysis of the amino acid sequence surrounding these O-linked glycosylation sites revealed that Pro was favoured at the +3 and –1 positions. This was consistent with previous reports, which gives some validation of the O-linked glycosylation sites identified using EXoO^35^, ^36^. However, achieving a better understanding the structural and functional roles of the O-linked glycans in these proteins certainly merits future investigation.

Compared to previous methods^2^, ^5^, ^10^, ^13^, ^21^, ^24^, EXoO identified a substantially larger number of O-linked glycosylation sites and glycoproteins almost doubling of the number of sites identified in recent decades. It also identified aberrant expression of 56 O-linked glycoproteins in kidney tumor tissue compared to normal tissue pointing to its utility in clinical investigations. Given these advantages of EXoO, it is anticipated that it will be widely applied in studies to analyse O-linked glycosylation of proteins.

## Methods

### Solid-phase extraction of site-specific O-linked glycopeptides from fetuin

Bovine fetuin (P12763) were denatured in buffer containing 8 M urea and 1 M ammonium bicarbonate (AB) and reduced in 5 mM DTT at 37°C for 1 hour. Proteins were alkylated in 10 mM iodoacetamide at room temperature (RT) for 40 min in the dark. The resulting samples were diluted eight-fold using 100 mM AB buffer before adding trypsin (enzyme/protein ratio of 1/40 w/w) and incubating at 37°C for 16 hours. Following digestion, peptides were de-salted using a C18 column (Waters, Milford, MA) according to manufacturer’s instructions.

Peptides were conjugated to AminoLink resin (Pierce, Rockford, IL) as previously described^20^. Briefly, the pH of the peptide containing eluate of the C18 column was adjusted to 7.4 by adding phosphate buffer (pH 8.0). Peptides were then incubated with the resin (100 µg/100 µl resin, 50% slurry) and 50 mM sodium cyanoborohydride (NaCNBH_3_) at RT for at least 4 hours or overnight with shaking. The resin was then blocked by adding 1M Tris-HCl buffer (pH7.4) containing 50 mM NaCNBH_3_ at RT for 30 min. The resin was washed three times with 50% acetonitrile, 1.5 M NaCl, and 20 mM Tris-HCl buffer (pH 6.8). O-linked glycopeptides were released from the resin by incubation with OpeRATOR^™^ and SialEXO^™^ (1 unit/1 µg peptides each enzyme, Genovis Inc, Cambridge, MA) in 100 µl of 20 mM Tris-HCl buffer (pH 6.8) at 37°C for 16 hours according to manufacturer’s instructions. Following this 16-hour incubation, the released peptides in solution were collected, and the resin was washed twice with 20 mM Tris-HCl buffer (pH 7.4) to collect the remaining peptides. The pooled peptides were then desalted on a C18 column and dried by lyophilization.

### Extraction of O-linked glycopeptides from human kidney tissue, serum and T cells

Collection and use of human tissue has been approved by Johns Hopkins Institutional Review Board (IRB). Kidney tumors were categorised as being clear cell renal cell carcinomas (CCRCC) and samples of tumor tissue were stored at –80°C before use. Control ormal kidney tissue samples were collected from the same individuals. Proteins from human kidney tissues, serum (Sigma-Aldrich, St. Louis, MO) and CEM T cells were trypsin-digested as described above. Following digestion, guanidination of peptides was conducted on a C18 column using procedure described previously to recover the Lys-containing peptides from complex samples^17^. Briefly, peptides were loaded on a pre-conditioned C18 column. The column was then sequentially washed three times with 0.1% TFA and then guanidination solution (equal volumes of 2.85 M aqueous ammonia, 0.6 M O-methylisourea and 0.1% TFA, final pH 10.5). After the final wash, guanidination solution was added to cover the C18 material in the column and it was incubated at 65°C for 20 min. Following this incubation, the column was cooled to RT and washed three time with 0.1% TFA. Peptides were eluted in 60% acetonitrile/0.1% TFA. Intact glycopeptides were enriched using a SAX HyperSep^™^ Retain AX Columns (RAX) column^9^. Briefly, after C18 desalting peptides in 60% acetonitrile/0.1% TFA were adjusted to 95% acetonitrile/1% TFA. The RAX column was conditioned in acetonitrile, 100 mM triethylammonium acetate, water and finally 95% acetonitrile/1% TFA using three times per solution. Samples were loaded, washed three times using 95% acetonitrile/1% TFA and eluted in 50% acetonitrile/0.1% TFA. The samples were then dried by lyophilization.

### Peptide fractionation

Peptides (100 µg) were split into 96 fractions using a 1220 Series HPLC (Agilent Technologies, Inc., CA) equipped with a Zorbax Extend-C18 analytical column containing 1.8 μm particles at a flow rate of 0.3 ml/min. The mobile-phase A was 10 mM ammonium formate (pH 10) and B was 10 mM ammonium formate and 90% acetonitrile (pH10). Peptides were separated using the following linear gradient: 0–2% B, 10 min; 2–8% B, 5 min; 8–35% B, 85 min; 35–95% B, 5 min; 95–95% B, 15 min. Fractions were collected from 0 to 96 min. The 96 fractions were concatenated into 24 fractions. The samples were then dried by lyophilization.

### LC-MS/MS analysis

Peptides dissolved in 0.1% formic acid (FA) were analyzed on a Fusion Lumos mass spectrometer with an EASY-nLC 1200 system or a Q-Exactive HF mass spectrometer (Thermo Fisher Scientific, Bremen, Germany) with a Waters NanoAcquity UPLC (Waters, Milford, MA). The mobile phase flow rate was 0.2 μL/min with 0.1% FA/3% acetonitrile in water (A) and 0.1% FA/90% acetonitrile (B). The gradient profile was set as follows: 6% B for 1 min, 6−30% B for 84 min, 30−60% B for 9 min, 60−90% B for 1 min, 90% B for 5 min and equilibrated in 50% B, flow rate was 0.5 μL/min for 10 min. MS analysis was performed using a spray voltage of 1.8 kV. Spectra (AGC target 4 × 10^5^ and maximum injection time 50 ms) were collected from 350 to 1800 m/z at a resolution of 60 K followed by data-dependent HCD MS/MS (at a resolution of 50 K, collision energy 29, intensity threshold of 2 × 10^5^ and maximum IT 250 ms) of the 15 most abundant ions using an isolation window of 0.7 m/z. Charge-state screening was enabled to reject unassigned, single, and more than six protonated ions. Fixed first mass was 110 m/z. A dynamic exclusion time of 45s was used to discriminate against previously selected ions.

### Database search of site-specific O-linked glycopeptides

Bovine fetuin (P12763) in a database with HIV gp120 (AAB50262.1) and TGFbeta1 (P01137) were used for the analysis of fetuin using the same procedure for search of the human protein database except that all fetuin peptides and four glycans including Hex(1)HexNAc(1), HexNAc, Hex(1)HexNAc(2) and Hex(2)HexNAc(2) were used. The RefSeq human protein database (72,956 sequences, downloaded from NCBI website Mar 25, 2015) was used to generate a randomized decoy database (decoy at protein level: decoy-pro) using The Trans-Proteomic Pipeline (TPP)^37^. The target and decoy protein database were concatenated and digested on the C-terminal side of Lys/Arg with 2 miss-cleavage sites (trypsin digestion) followed by digested on the N-terminal side of Ser/Thr with 5 miss-cleavage sites (OpeRATOR digestion) in silico and finally Ser/Thr containing peptides with peptide lengths between 6 to 46 amino acids were used resulting in 30,759,520 non-redundant peptide entries. SEQUEST in Proteome Discoverer 2.2 (Thermo Fisher Scientific) was used to search against the database with oxidation (M), guanidination (K), Hex(1)HexNAc(1) (S/T) and HexNAc (S/T) as the variable modifications. Static modification was carbamidomethylation (C). FDR set at 1% using Percolator. MS/MS scan numbers of oxonium ion containing spectra were extracted with 10 ppm tolerance from raw files. Oxonium ion 204 was mandatory together with two of the other oxonium ions. The result was filtered to report identification with glycan modification and oxonium ions in the MS/MS spectra. Percolator generated FDR was verified using peptide identification labelled with decoy-pro in the output result.

### Label-free quantification using spectral counting

PSM numbers with 1% FDR were counted for peptides identified from tumor or normal tissue. The data was then normalized using the total number of PSMs in individual sample. The fold change of peptides between tumor and normal samples was calculated. Peptides presented exclusively in tumor or normal tissue samples were given a 2- or 0.5-fold change, respectively, in order that a fold change at the protein level could be calculated. The average fold change of peptides was calculated to obtain a fold change of proteins and at least 2-fold change for proteins was used. In addition, the otal number of PSMs for proteins in samples was summed and a difference of at least 10 PSM between tumor derived and normal tissue samples was used to determine significant change.

### Bioinformatics

Site-specific O-linked glycopeptides were re-constructed into peptides of 15 amino acids in length with the O-linked glycosylation sites as the central amino acid. Online software pLogo was used to predict the motifs^38^. Hierarchical clustering was used to cluster the samples based on the number of PSM of site-containing peptides using QCanvas version 1.21^39^. The Database for Annotation, Visualization and Integrated Discovery (DAVID) and UniProt (http://www.uniprot.org) were used for Gene Ontology (GO) analysis^40^.

### Accession code

The LC-MS/MS data have been deposited to the PRIDE partner repository^41^ with the dataset identifier: project accession: PXD009476.

Reviewer account details:

Username

Password: hFtDLRPy

## Acknowledgement

We acknowledge Prof. Malcolm McCrae for proof-reading and Dr. David Clark for maintenance of Fusion Lumos Mass Spectrometer. We acknowledge service from Johns Hopkins Mass Spectrometry and Proteomics Core. This work was supported by the National Institutes of Health, the National Institute of Allergy and Infectious Diseases (R21AI122382), National Cancer Institute, the Early Detection Research Network (EDRN, U01CA152813), the Clinical Proteomic Tumor Analysis Consortium (CPTAC, U24CA210985), National Heart Lung and Blood Institute, Programs of Excellence in Glycosciences (PEG, P01HL107153), and by amfAR, The Foundation for AIDS Research on Bringing Bioengineers to Cure HIV (Grant amfAR 109551–61-RGRL).

## Contributions

W.Y. and H.Z. conceived and wrote the manuscript; W.Y. conducted experimental and data analysis; W.Y. and M.A. conducted programming, data analysis, statistics and bioinformatics;

W.Y and Y.H. developed strategy for identification of glycopeptides; Q.K.L. collected and conducted pathological examination for tumor and normal kidney tissues.

---

**Supplementary Figure 1.**
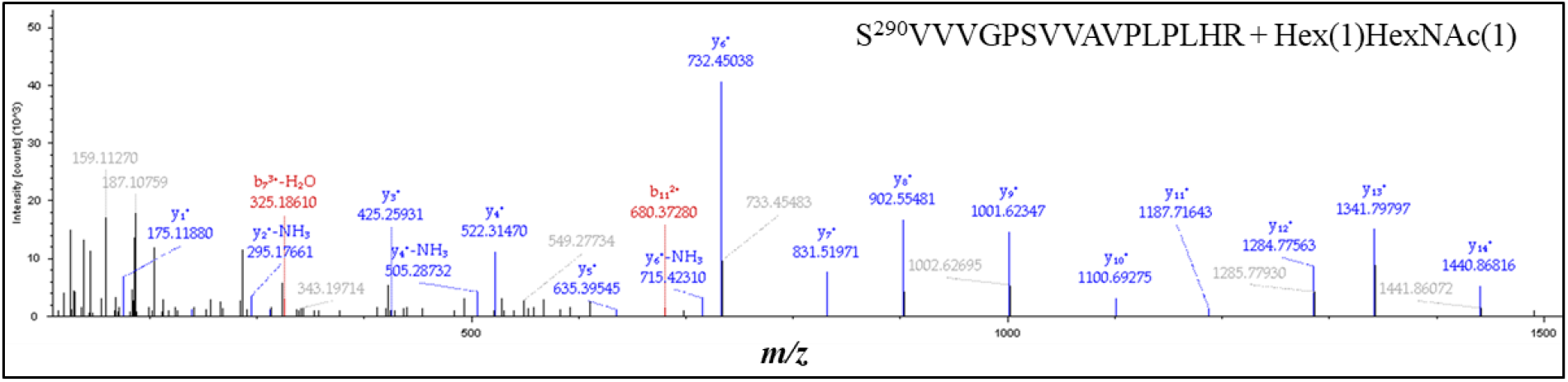
Identification of a new O-linked glycosylation site at Ser-290 in bovine fetuin.

**Supplementary Figure 2.**
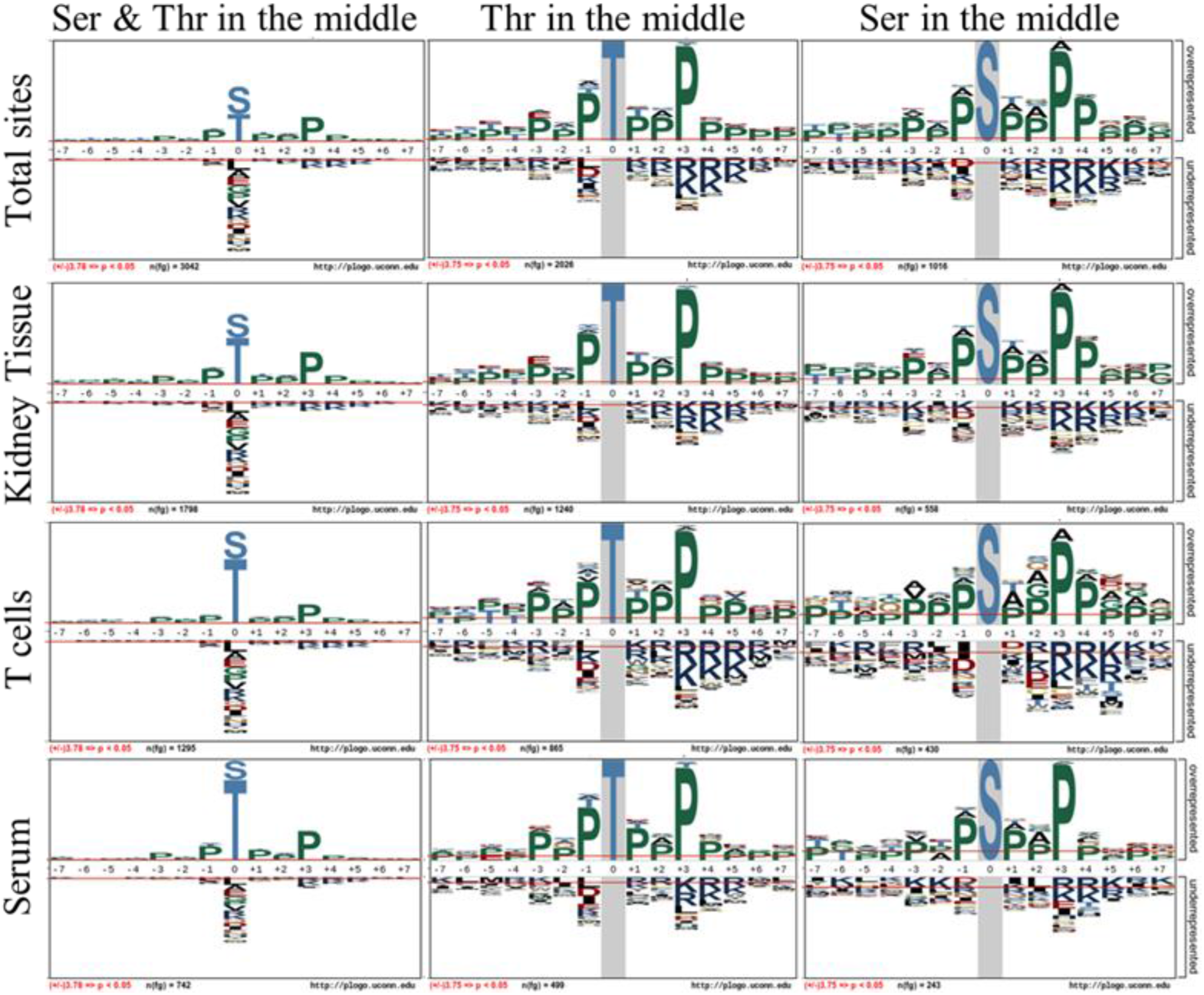
Motif analysis of the O-linked glycosylation sites in different samples.

**Supplementary Figure 3.**
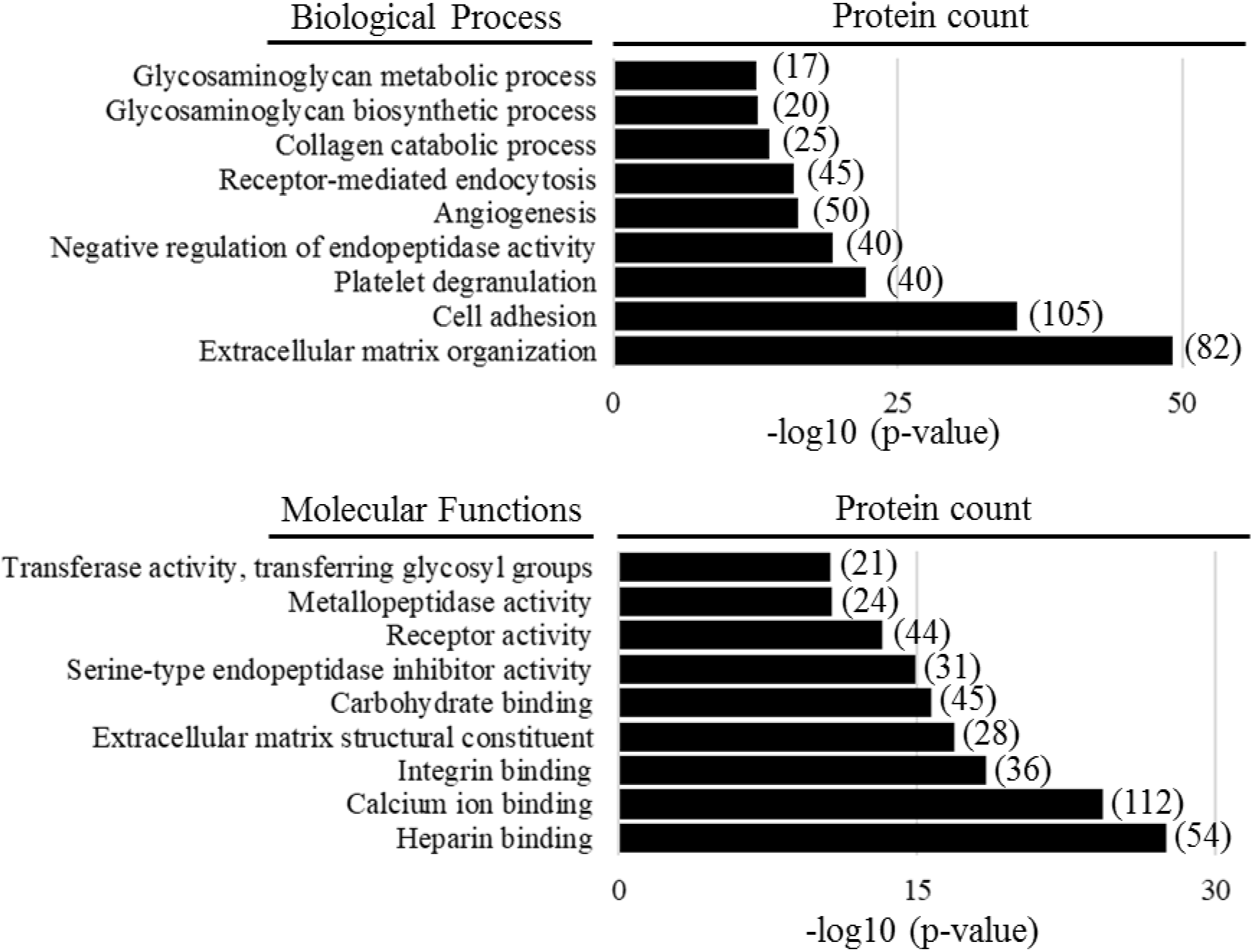
Biological process and molecular functions of the identified O-linked glycoproteins.

**Supplementary Figure 4.**
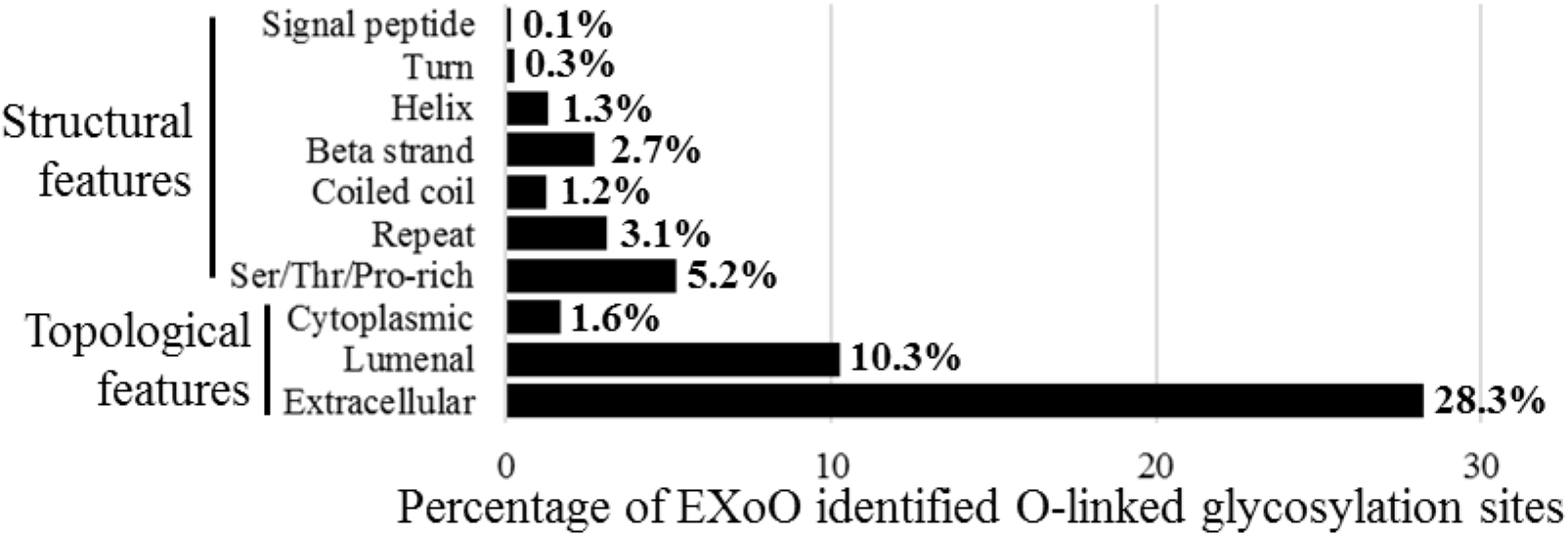
Localization of sites to protein structural and topological features.

**Supplementary table 6.** Differentially expressed O-linked glycoproteins in tumor vs normal kidney tissue.

| Protein Accession # | Gene Name | Protein Name | Fold Change tumor/normal (log2) | Differential PSM between tumor and normal |
| --- | --- | --- | --- | --- |
| P05121 | SERPINE1 | Plasminogen activator inhibitor 1 | Tumor | 18 |
| Q16790 | CA9 | Carbonic anhydrase 9 | 3.7 | 12 |
| Q8NHJ6 | LILRB4 | Leukocyte immunoglobulin-like receptor subfamily B member 4 | 3.0 | 14 |
| P27658 | COL8A1 | Collagen alpha-1(VIII) chain | 2.7 | 127 |
| P48960 | CD97 | EGF-like module-containing mucin-like hormone receptor-like 2 | 2.5 | 14 |
| P19823 | ITIH2 | Inter-alpha-trypsin inhibitor heavy chain H2 | 2.5 | 415 |
| P50281 | MMP14 | Matrix metalloproteinase-14 | 2.5 | 18 |
| Q68BL8 | OLFML2B | Olfactomedin-like protein 2B isoform 2 | 2.3 | 15 |
| Q99715 | COL12A1 | Collagen alpha-1(XII) chain | 2.3 | 412 |
| P15502 | ELN | Elastin | 2.2 | 13 |
| P49257 | LMAN1 | Protein ERGIC-53 | 2.2 | 14 |
| P98095 | FBLN2 | Fibulin-2 | 2.1 | 247 |
| Q14767 | LTBP2 | Latent-transforming growth factor beta-binding protein 2 | 2.0 | 198 |
| Q96DB9 | FXYD5 | FXYD domain-containing ion transport regulator 5 | 1.9 | 17 |
| Q9BY76 | ANGPTL4 | Angiopoietin-related protein 4 | 1.8 | 96 |
| Q9UKP4 | ADAMTS7 | A disintegrin and metalloproteinase with thrombospondin motifs 7 | 1.7 | 88 |
| P06213 | INSR | Insulin receptor | 1.6 | 13 |
| Q12805 | EFEMP1 | EGF-containing fibulin-like extracellular matrix protein 1 | 1.6 | 258 |
| Q92896 | GLG1 | Golgi apparatus protein 1 | 1.5 | 12 |
| Q9BXX0 | EMILIN2 | EMILIN-2 | 1.5 | 58 |
| P16112 | ACAN | Aggrecan core protein | 1.5 | 832 |
| Q9UBP4 | DKK3 | Dickkopf-related protein 3 | 1.4 | 32 |
| Q9NS15 | LTBP3 | Latent-transforming growth factor beta-binding protein 3 | 1.4 | 12 |
| Q12797 | ASPH | Aspartyl/asparaginyl beta-hydroxylase | 1.3 | 42 |
| P23470 | PTPRG | Receptor-type tyrosine-protein phosphatase gamma | 1.3 | 12 |
| Q9H8L6 | MMRN2 | Multimerin-2 | 1.3 | 14 |
| P13611 | VCAN | Versican core protein | 1.3 | 1216 |
| Q8IUX7 | AEBP1 | Adipocyte enhancer-binding protein 1 | 1.2 | 42 |
| Q8TDQ1 | CD300LF | CMRF35-like molecule 1 | 1.2 | 11 |
| Q14515 | SPARCL1 | SPARC-like protein 1 | 1.1 | 40 |
| Q14766 | LTBP1 | Latent-transforming growth factor beta-binding protein 1 | 1.1 | 13 |
| Q14314 | FGL2 | Fibroleukin | 1.1 | 27 |
| Q6UXD5 | SEZ6L2 | Seizure 6-like protein 2 | 1.1 | 19 |
| P02751 | FN1 | Fibronectin | 1.1 | 218 |
| P16070 | CD44 | CD44 antigen | 1.0 | 17 |
| Q6IEE7 | TMEM132E | Transmembrane protein 132E | -1.0 | 14 |
| P15144 | ANPEP | Aminopeptidase N | -1.1 | 181 |
| O60494 | CUBN | Cubilin | -1.2 | 66 |
| Q8N474 | SFRP1 | Secreted frizzled-related protein 1 | -1.2 | 47 |
| O75309 | CDH16 | Cadherin-16 | -1.2 | 14 |
| Q9NR34 | MAN1C1 | Mannosyl-oligosaccharide 1,2-alpha-mannosidase IC | -1.2 | 11 |
| P98164 | LRP2 | Low-density lipoprotein receptor-related protein 2 | -1.2 | 818 |
| O43451 | MGAM | Maltase-glucoamylase | -1.3 | 62 |
| P02724 | GYPA | Glycophorin-A | -1.3 | 40 |
| Q9ULC0 | EMCN | Endomucin | -1.4 | 142 |
| O95428 | PAPLN | Papilin | -1.4 | 79 |
| Q9UGT4 | SUSD2 | Sushi domain-containing protein 2 | -1.6 | 14 |
| Q9UJW2 | TINAG | Tubulointerstitial nephritis antigen | -1.6 | 60 |
| Q6ZMM2 | ADAMTSL5 | ADAMTS-like protein 5 | -1.7 | 13 |
| P36269 | GGT5 | Gamma-glutamyltransferase 5 | -1.7 | 13 |
| Q5IJ48 | CRB2 | Protein crumbs homolog 2 | -1.9 | 22 |
| Q96SM3 | CPXM1 | Probable carboxypeptidase X1 | -2.0 | 13 |
| Q96A83 | COL26A1 | Collagen alpha-1(XXVI) chain | -2.1 | 23 |
| Q2UY09 | COL28A1 | Collagen alpha-1(XXVIII) chain | Normal | 15 |
| P13866 | SLC5A1 | Sodium/glucose cotransporter 1 | Normal | 20 |
| Q8N695 | SLC5A8 | Sodium-coupled monocarboxylate transporter 1 | Normal | 26 |

